# Direct Reciprocity Under Uncertainty Does Not Explain One-shot Cooperation, but Demonstrates the Benefits of a Norm Psychology

**DOI:** 10.1101/004135

**Authors:** Matthew R. Zefferman

**Affiliations:** Graduate Group in Ecology, University of California, Davis, U.S.A National Institute for Mathematical and Biological Synthesis, University of Tennessee, U.S.A.

## Abstract

Humans in many societies cooperate in economic experiments at much higher levels than would be expected if their goal was maximizing economic returns, even when their interactions are anonymous and one-shot. This is a puzzle because paying a cost to benefit another in one-shot interactions gives no direct or indirect benefits to the cooperator. This paper explores the logic of two competing evolutionary hypotheses to explain this behavior. The “norm psychology” hypothesis holds that a player's choice of strategy reflects socially-learned cultural norms. Its premise is that over the course of human evolutionary history, cultural norms varied considerably across human societies and through a process of gene-culture co-evolution, humans evolved mechanisms to learn and adopt the norms that are successful in their particular society. The “mismatch” hypothesis holds that pro-social preferences evolved genetically in our hunter-gatherer past where one-shot anonymous interactions were rare and these preferences are misapplied in modern laboratory settings. I compare these hypotheses by adopting a well-known model of the mismatch hypothesis and show that selection for one-shot cooperation in the model is an artifact of agents being constrained to only two strategies: Tit-for-Tat and Always Defect. Allowing for repentant and forgiving strategies reverses selection away from one-shot cooperation under all environmental parameters. Direct reciprocity does not necessarily lead to cooperation, but instead generates many different normative equilibria depending on a group’s idiosyncratic evolutionary history. Therefore, an agent whose behavior is evoked solely from non-cultural environmental cues will be disadvantaged relative to an agent who learns the locally successful norms. Cooperation in one-shot laboratory experiments is thus more easily explained as the result of a psychology evolved for learning social norms than as a genetic mismatch.

## 1. Introduction

A puzzling finding in experimental economics is that in many societies humans cooperate in laboratory experiments at much higher levels than they would if they were money-maximizing agents. Players act as though they have “pro-social preferences,” that is, in addition to their own welfare, they care about some combination of the welfare of other players, fairness and equality (Fehr and Schmidt, 1999; Bolton and Ockenfels, 2000). The strongest evidence for pro-social preferences come from simple games, such as the “dictator game,” where participants are given an amount of money that they can divide between themselves and an anonymous stranger. While the money-maximizing agent would keep the entire sum, players consistently distribute substantial sums to anonymous strangers (Camerer, 2003). Pro-social play has also been observed in more complicated games, such as ultimatum games, trust games, Prisoner’s Dilemmas, and public goods games (Fehr and Schmidt, 1999; Bolton and Ockenfels, 2000; Camerer, 2003) and has been documented in many societies, though there there is substantial variation both within and between societies (Henrich et al., 2004). Pro-social play is especially puzzling when a game is played only once with an anonymous partner since one-shot anonymous interactions eliminate reciprocity and reputation-building as potential motivations. What explains the existence of pro-social play in one-shot anonymous games?

In this paper I explore two competing evolutionary hypotheses for the origins of pro-social play in one-shot economic experiments. One, the “norm psychology” hypothesis, posits that cooperative play reflects cultural norms acquired during a player’s lifetime through social learning (Richerson and Boyd, 2005; Henrich and Henrich, 2007; Boyd and Richerson, 2009; Chudek and Henrich, 2011; Chudek et al., 2013). This hypothesis is premised on the proposition that humans tend to adopt the norms of their group and over the course of humans’ evolutionary history, the norms of different human groups, even in similar environments, were highly varied and subject to frequent change. Variation in norms between societies was partly due to variation in local ecology. However, theoretical models suggest that, even when ecological circumstances are held constant, cultural change is path dependent. The combination of repeated interaction, punishment, and conformity can stabilize, at least temporarily, a wide range of behaviors (Boyd and Richerson, 1992; Boyd, 2006; Henrich and Henrich, 2007). Because successful behaviors for one’s particular society would be difficult to infer from the local ecology alone and because cultural change is often much faster than genetic selection, humans evolved a “norm psychology” that helps us learn and adopt the currently prevailing norms of our *particular* society.

Because different groups, even in similar environments, are expected to develop different norms, the norm psychology hypothesis predicts that there will be substantial cultural variation between groups that can be operated on by cultural selection. Groups with more cooperative norms will tend to out-compete others (Boyd and Richerson, 1990). However, since not every group will be maximally cooperative as individuals within groups are tempted to shirk from cooperative endeavors for personal gain, there will still be variation between groups in norms for cooperation in social exchange. In economic experiments, players’ cooperation will reflect the prevailing norms of their particular society. This helps explain between-society variation in levels of cooperation in economic experiments (Henrich et al., 2004).

A competing hypothesis is a type of “mismatch” hypothesis (Price, 2008; Chudek et al., 2013) called, variously, the “Savanna Principle” (Kanazawa, 2004), the “big mistake hypothesis” (Richerson and Boyd, 2005), the “misapprehension hypothesis” (Hagen and Hammerstein, 2006), the “evolutionary legacy hypothesis”(Burnham and Johnson, 2005) and “social exchange theory” (Krasnow et al., 2012). The premise of this hypothesis is that cooperative play in one-shot experiments is primarily due to adaptations acquired in ancient human environments through selection on genes (Kanazawa, 2004; Hagen and Hammerstein, 2006; Price, 2008; Delton et al., 2011a; Pinker, 2012; Krasnow et al., 2012; McCullough et al., 2013; Pedersen et al., 2013).

The mismatch hypothesis posits that over the course of humans’ evolutionary history, there were very few one-shot or anonymous encounters since “observations and the demographic conditions that characterize hunter-gatherer life indicate that large numbers of repeat encounters, often extending over decades, was a stable feature of the social ecology of ancestral humans” (Delton et al., 2011a). Although these “theories predict that our evolved social psychology will be calibrated by relevant environmental inputs” (Delton et al., 2010), this calibration is imperfect because genetic selection in ancient environments did not adequately prepare humans for the types of anonymous interactions that are more common in modern society (including laboratory experiments). Furthermore, because one-shot interactions were so rare in our ancient past, humans who were endowed with a propensity to cooperate in an initial interaction would do better because they would more easily capture the benefits of future interactions that would most likely follow. Instead of reflecting socially-learned and local norms, one-shot cooperation in laboratory experiments “is the expected expression of evolutionarily well-engineered decision-making circuitry specialized for effective reciprocity” (Delton et al., 2011a). There is, thus, an “evolutionary mismatch” between genetically-evolved decision-making circuitry and modern environments or, in other words, our “modern skulls housing a stone age mind” (Cosmides and Tooby, 1997).

Table 1 summarizes key differences between the norm psychology and mismatch hypotheses for explaining one-shot cooperation in economic experiments. While both hypotheses rely on genetically evolved psychology, the role of this psychology is different. In the norm psychology hypothesis, it provides rules for adopting local norms (such as who to copy), including norms for social exchange. In the mismatch hypothesis, evolved psychology provides rules for social exchange directly through genetically-evolved “protocols” (Price, 2008) or “decision-making circuitry” (Delton et al., 2011a). The hypotheses also differ as to what inputs are important for influencing human behavior. Both hypotheses accept that certain environmental inputs, such as the benefits to cooperation and the probability of repeated interactions inform players’ strategy choices (e.g., Delton et al., 2010; Henrich and Henrich, 2007, 56). However, while the norm psychology hypothesis stipulates individuals rely to large extent on mechanisms for socially-learning strategies that will be successful in their particular group (Henrich and Henrich, 2007, 47-59), the mismatch hypothesis explicitly denies the need to invoke such learning to explain cooperative behavior (Burnham and Johnson, 2005; Price, 2008; Delton et al., 2011a). Finally, the hypotheses differ in how they view the role of repeated interaction in human evolutionary history. Proponents of the mismatch hypothesis predict that if interactions are sufficiently repeated, especially if the benefits to cooperation are high, genetic selection for cooperation under these conditions would be inevitable (e.g., Delton et al., 2011a; McCullough et al., 2013). Proponents of the norm psychology hypothesis predict that even when the potential benefits to cooperation are high and, especially, when interactions are sufficiently repeated, there can be great variation in the amount of cooperation that might evolve, culturally, in a given group (e.g., Henrich and Henrich, 2007).

**Table 1.**
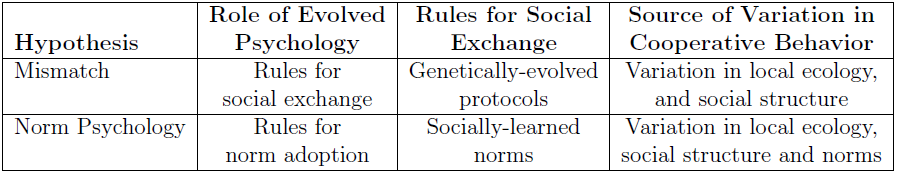
Key differences between the mismatch and norm psychology and hypotheses

Fehr and Henrich (2003), Henrich et al. (2004) and Chudek et al. (2013) have raised empirical objections to the mismatch hypothesis based primarily on ethnographic and experimental evidence. They point to ethnographic evidence that hunter-gatherers have many more interactions with strangers than is supposed by proponents of the mismatch hypothesis. They also point to evidence that participants in economic experiments seem quite capable of distinguishing one-shot from repeated games and, in fact, adjust their behavior accordingly. However, others have found these objections unconvincing (Hagen and Hammerstein, 2006) and question the plausibility of the norm psychology hypothesis as an alternative (Price, 2008; Delton et al., 2010). This empirical debate is important, but beyond the scope of this paper. Instead, I focus on which hypothesis is better supported by theory. Towards this end, I adopt a model developed to formally explain and support the mismatch hypothesis (Delton et al., 2011a), showing that if certain artificial constraints on the scope of behaviors available to selection are relaxed, it provides better support for the norm psychology hypothesis.

### 1.1. The DKCT Model

I adopt a recent model by Delton, Krasnow, Cosmides and Tooby (2011) (hereafter “DKCT” and “the DKCT model”) which has sought to put the mismatch hypothesis on more solid theoretical footing. Like other proponents of the mismatch hypothesis, DKCT hypothesize that cooperation in one-shot economic experiments can be explained by genetic selection among “our band-living hunter-gatherer ancestors” (Delton et al., 2011a, S1) whose lives were dominated by repeated interactions. However, they add a twist. When strangers meet there is *uncertainty* about whether it will be a one-time encounter or a repeated interaction. Since repeated interaction can create, over time, greater absolute costs and benefits than one-shot interactions, mistaking a repeated interaction for a one-shot interaction is more costly than mistaking a one-shot interaction for a repeated one (see also Krasnow et al. (2012), Pinker (2012), McCullough et al. (2013), and Pedersen et al. (2013)). DKCT run a series of simulations of this idea and find that, as premised by the mismatch hypothesis, agents will evolve a propensity to cooperate, even if there is strong evidence that an interaction is one-shot. In this section I briefly describe the DKCT model, which is further elaborated in the supplemental materials of their paper (Delton et al., 2011a). I then describe reasons one might be skeptical of their results based on previous theory and show how the high levels of cooperation in one-shot interactions they find are an artifact of constraining their agents to an evolutionary history that allows only two of an infinite number of possible strategies. Then I show how groups exposed to slightly different evolutionary histories under the same environmental conditions will evolve very different patterns of behavior, as premised by the norm psychology hypothesis. Wherever I was unsure of the original model’s details, I consulted the authors who helpfully provided clarification.

In the DKCT model, 500 agents are born, randomly pair into dyads, play a game, reproduce, and die. They play either a one-shot or repeated Prisoner’s Dilemma (PD), where, in each round, they can pay a cost, *c*, to confer a benefit, *b*, to their partner (Fig. 1). With a probability, *P*, the game is one-shot. Otherwise it is repeated. If the game is repeated, after each round the probability that the game lasts another round is *w*. Therefore, given a repeated game, the number of rounds is drawn from a geometric distribution with an expectation of 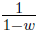.

**Figure 1.**
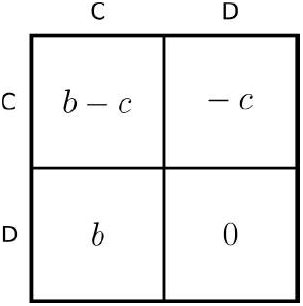
The payoffs for Player 1 in one round of a repeated Prisoner’s Dilemma game. An agent who cooperates (plays “C”) pays a cost c to provide a benefit b to its partner. An agent who defects (plays “D”) does not. Since defecting always gives a higher payoff than cooperating, a payoff-maximizing agent would always defect in a one-shot game. In a repeated game, the same pair of agents play this game multiple times.

Before playing the game each agent in a dyad independently receives an imperfect signal about whether the interaction is likely to be repeated or one-shot. As shown in Fig. 2, if the game is repeated, the signal is drawn from a normal distribution with a mean of *d*/2 and a standard deviation of one. If the game is one-shot, the signal is drawn from a normal distribution with a mean of −*d*/2 and a standard deviation of one. Since these distributions overlap, an agent cannot be sure which distribution the signal was drawn from. Since the amount of overlap decreases with the size of *d*, this parameter is a measure of the certainty in the model.

**Figure 2.**
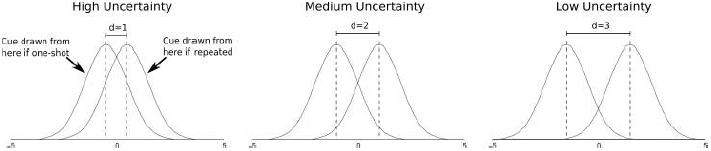
Nature draws a signal from one of two distributions depending on whether an interaction is one-shot or repeated. If a game is repeated, the signal is drawn from a normal distribution with a mean of *d*/2 and a standard deviation of one. Otherwise the signal is drawn from a distribution with a mean of −*d*/2. DKCT use three values of *d* with lower values indicating more uncertainty since there is more overlap between distributions.

Each agent is born with a “cue threshold,” a number it uses to pick a strategy based on the imperfect signal it receives. If an agent’s signal is greater than its cue threshold (indicating a repeated game), the agent plays Tit-for-Tat (TFT), a strategy that cooperates on the first round of play and thereafter repeats the actions of the other agent on the previous round. If an agent’s signal is less than its cue threshold (indicating a one-shot game), the agent plays Always Defect (ALLD), a strategy that defects on every round.

In the first generation, cue thresholds are distributed normally with a mean of 0 and a standard deviation 0.025. When an agent is born in later generations, it inherits a cue threshold from a member of the previous generation, with a probability proportional to the members’ relative payoffs. However there is a high, 5%, chance that an agent’s decision threshold will mutate, changing by a normally distributed random variable with a mean of 0 and a standard deviation 0.025. After reproducing, all members of the previous generation are removed from the population. DKCT’s simulations each lasted for 10,000 generations. Agents start each generation with a baseline payoff of 10.

DKCT run their simulations under 750 separate parameter conditions which are given in Table 2. For each parameter condition, DKCT measured the population’s expected probability of one-shot cooperation averaged over the last 500 generations, where the “expected probability of one-shot cooperation” is the expectation of a randomly-selected agent’s probability of cooperating (playing TFT) given that an interaction is one-shot. DKCT found that agents’ cue thresholds evolved to be low so that they had high probabilities of one-shot cooperation for many of these 750 parameter combinations. In fact, agents often developed a bias towards playing TFT even when there was strong evidence that an interaction was one-shot.^1^ However, the agents in DKCT’s simulations are confined to only two strategies, TFT and ALLD. As I describe below, a more complete set of possible strategies would generate a greater diversity of behavioral equilibria. I will show that if other strategies are allowed to compete with TFT in DKCT’s model, selection can act against one-shot cooperation until it virtually disappears from the population, demonstrating the that repeated interaction does not inevitably lead to cooperative equilibria.

**Table 2.** Table of parameters. DKCT and I both run our simulations under every combination of these parameters, for 750 total simulations. In addition, I run the model where each agent has only one partner in their lifetime (as in DKCT) and where each agent has ten lifetime partners.

| Symbol | Definition | Values |
| --- | --- | --- |
| $c$ | Cost of cooperating in a round | 1 |
| $b$ | Benefits from cooperation to other player in one round | 1.5, 2, 3, 4, 5, 6, 7, 8, 9, 10 |
| $P$ | Probability that an interaction is one-shot | 0.1, 0.3, 0.5, 0.7, 0.9 |
| $w$ | In a repeated interaction, probability of a subsequent round | 0.5, 0.7, 0.9, 0.95, 0.99 |
| $d$ | Amount of uncertainty over whether a game is one-shot or repeated | 1, 2, 3 |

### 1.2. Repeated Interaction Creates Many Unstable Equilibria

Repeated games do not necessarily favor cooperation, but have many behavioral equilibria with cooperation ranging anywhere from 0% to 100% (Bendor and Swistak, 1997), with the specific equilibrium dependent on the distribution of strategies in the population and the order in which novel strategies invade. DKCT’s simulations use only TFT and ALLD. They chose TFT, in part, because it had the “benefit of being familiar to most readers.” TFT is familiar to most readers because it famously performed better than any other strategy submitted to two computerized tournaments (Axelrod and Hamilton, 1981; Axelrod, 1984). TFT was one of the simplest strategies in the tournaments and since it was both “nice” (cooperating on the first turn) and “retaliatory” (defecting after a partner defects), its success seemed to cement “niceness” and “retaliation” as the paths to evolutionary success in repeated games.

However, TFT’s success hinged on the particular mix of strategies entered in the tournaments and does not generalize to other mixes of strategies. One reason is explained by the “folk theorem” of repeated games. Well-known to game theorists by the 1950s and 1960s, it shows that if a PD is sufficiently repeated, *any* pattern of behavior can be an equilibrium (Rubinstein, 1979; Fudenberg and Maskin, 1986). All that is required is that players adhere to a pattern and retaliate against any other player that deviates from the pattern. This is a problem for the premise that repeated interactions inevitably lead to cooperative equilibria, as both cooperative and non-cooperative equilibria are possible. As Boyd (2006) puts it, “when everything is an equilibrium, showing that reciprocity is an equilibrium too does not really tell you much.”

Another reason why cooperation is not the inevitable outcome of repeated PDs is that none of these equilibria are evolutionarily stable. They can *all* be invaded by other strategies. This was quickly pointed out in the case of TFT (Williams, 1984; Selton and Hammerstein, 1984), after Axelrod and Hamilton (1981)’s initial claims of its stability. It was later proved that there are *no* evolutionarily stable strategies in a sufficiently repeated PD (Boyd and Lorberbaum, 1987; Farrell and Ware, 1989; Lorberbaum, 1994). For example, Boyd and Lorberbaum (1987) show that TFT can be invaded by a more forgiving strategy called “Tit-for-Two Tats” (TF2T) that is nice but only retaliates after its opponent defects twice in a row, in combination with a nasty strategy called “Suspicious Tit-for-Tat”(STFT) that is similar to TFT except that it is “nasty,” defecting on the first turn. Agents playing TF2T increase in the population relative to TFT since agents playing it have higher payoffs when playing STFT. However, agents with a strong bias towards playing STFT exploit TF2T’s forgiveness and eventually invade to near fixation. This is an example of selection in repeated interactions destabilizing a nice strategy, like TFT, and promoting a nasty strategy, STFT. By limiting their simulations to TFT and ALLD, DKCT do not allow for this type of invasion by nasty strategies.

This type of invasion also occurs in computational models. For example, Axelrod (1997, 17-22) used genetic algorithms to evolve strategies to play against a representative sample of the strategies submitted to his tournament. While TFT-like strategies evolved in these simulations, the very best strategies (i.e., those that performed even better than TFT) all defected on the first move. According to Axelrod, these high-performing strategies “had responses that allowed them to ‘apologize’ and get to mutual cooperation with most of the unexploitable representatives and different responses that allowed them to exploit a representative that was exploitable.” Similarly, Nowak and Sigmund (1989) showed that, even in a set of fairly simple strategies, the evolutionary dynamics of agent’s playing iterated games are more complex than “cooperation wins.” For example, even without changing payoff structures or other exogenous parameters, selection can cycle indefinitely between favoring relatively “nice” and relatively “nasty” strategies. In the next sections, I show that DKCT’s type of uncertainty does not make strategies with high probabilities of one-shot cooperation immune to invasion by nastier strategies under any of the parameters they model. I then describe why it is better for an agent to learn the norms prevalent in its society than it is to instinctively cooperate in a repeated interaction.

## 2. Methods

DKCT’s simulations start with a very specific proposition. Agents will always defect if given sufficient evidence that an interaction is one-shot and play TFT if given sufficient evidence than an interaction will be repeated. However, as explained above, theory indicates that any behavior is a plausible equilibrium in a repeated PD. DKCT’s model is different than a standard repeated PD because they include a degree of uncertainty over whether a game is one-shot or repeated. Does this change the logic of repeated games?

To answer this question, I replicated DKCT’s model with different combinations of initial strategies. To ensure that any differences between outcomes of these combinations were due only to the mix of strategies, I also replicated the aspects of DKCT’s model that were particularly friendly to generating cooperative outcomes. First, all interactions involve only two players, which is the condition where reciprocity most easily generates cooperation. In fact, reciprocity becomes *geometrically* less effective as more players are added to a game (Joshi, 1987; Boyd and Richerson, 1988). Second, their simulations *begin* with every agent playing a cooperative strategy with enough evidence that a game is repeated. This is a strong assumption because the more likely ancestral condition would have little to no altruistic cooperation and it is much harder for reciprocity to explain cooperation’s *origins* than its maintenance. Third, the parameter values were very friendly to cooperation. A single cooperative act could have up to a 1000% return on investment and this high return could potentially be realized over hundreds of interactions. To bias my simulations towards DKCT’s findings, I retain all of these cooperation-favoring assumptions.

I ran simulations of the DKCT model for all 750 of their parameter combinations under four initial conditions. To demonstrate consistency with DKCT’s original findings, the first condition is an exact replication of their simulations which include only TFT and ALLD. In the other three conditions, I introduce strategies not considered by DKCT. Since written descriptions of strategies for repeated games can sometimes be ambiguous, I precisely represent the strategies of all treatments, following Rubinstein (1986) and Miller (1996), as Finite State Automata in Appendix D.

I also ran each simulation under two conditions for the number of partners an agent has in its lifetime. In one condition, as in DKCT’s original simulations, each agent only interacts with one other agent. In the other, agents interact with ten others. This condition is not only more realistic, since most humans interact with multiple other people in their lifetime, but it also decreases stochastic shocks due to skew in the number of rounds each agent plays. A quirk of having only one interaction partner is that under some parameter conditions (i.e., high *P* and *w*) almost every dyad plays just one round, but a small subset plays in the hundreds. This creates shocks where low-performing strategies jump to near fixation simply because a dyad was randomly assigned a game with substantially more rounds than most other dyads combined. However, as I describe in Appendix C, when agents interact with multiple partners the distribution of rounds played by each agent is substantially less skewed. I present the results of both one-partner and ten-partner conditions in Appendix A.

### 2.1. Treatment 1: TFT and ALLD

In Treatment 1, as in DKCT’s model, agents play TFT if their signal is above their cue threshold (indicating a higher probability of a repeated game) and ALLD if the signal is below their threshold (indicating a higher probability of a one-shot game). Fig. 3A shows the expected payoffs for one-shot and repeated games for agents playing TFT and ALLD. I replicate their model for all 750 combinations of parameter values explored by DKCT and with agents having only one lifetime partner (as in DKCT) or ten lifetime partners.

**Figure 3.**
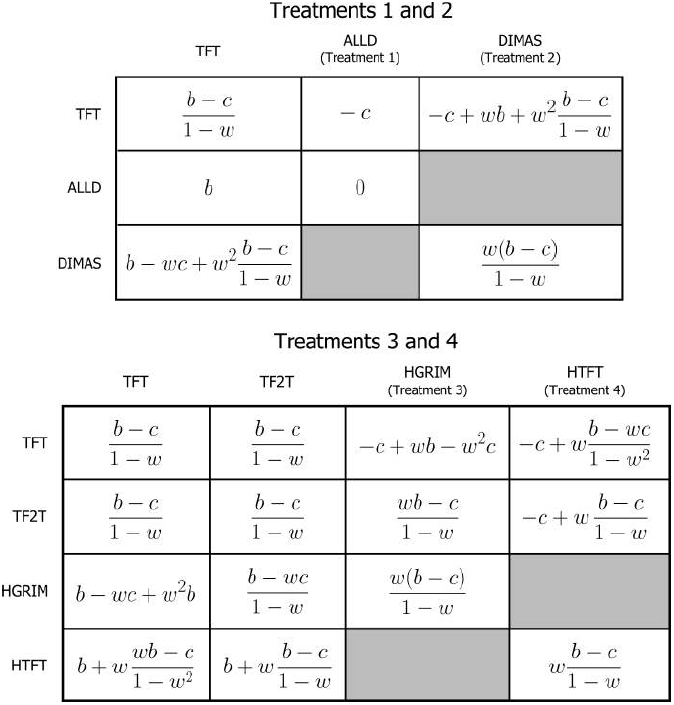
Expected payoffs for each strategy in the four treatments, assuming a repeated game. *b* is the benefits to cooperation, *c* is the cost of cooperating, and w is the probability that, after each round in a repeated game, there is another round. Note that the number of rounds in a repeated game is geometrically distributed where the expectation of the number of rounds is 1/(1 - *w*). **A** shows the expected payoffs for Treatments 1 and 2. **B** shows the expected payoffs for Treatments 3 and 4.

### 2.2. Treatment Two: TFT and DIMAS

In DKCT’s original simulations first-round defectors must continue, by assumption, to defect for all time. In other words, agents cannot repent or, in Axelrod (1997)’s terminology, “apologize.” However, the ability to make amends is an important part of most humans’ behavioral repertoires. McNally and Tanner (2011) suggest that a repentant strategy would perform better than ALLD in DKCT’s model, reducing the amount of one-shot cooperation. To test this suggestion, I replace ALLD with a simple strategy that defects on the first round, but finding itself in a repeated game, immediately repents and begins cooperating. I dub this strategy “DIMAS” after the biblical thief whose repentance earns him eternal rewards in paradise.

Figure 3B shows the expected payoffs for one-shot and repeated games for agents playing TFT when the signal is above their cue threshold and DIMAS when the signal is below it. These are similar to those in Fig. 3A except that DIMAS typically earns higher payoffs than ALLD because it both cooperates with itself starting in round two and cooperates with TFT starting in round three. DIMAS, with only two memory states (see Appendix D), is the simplest case of a large class of repentant strategies.

### 2.3. Treatment Three: TFT, TF2T and HGRIM

Delton et al. (2011b), in a response to McNally and Tanner (2011), state that for a model including both repentant and forgiving strategies, ”[w]e concur that the fitness differential between initial cooperation and defection might be smaller in such a model. Importantly, however, the direction of selection would be the same, just with reduced strength. Hence, the effects would not be eliminated and the generality of our results would be unchanged.” Their implication, with which I agree, is that the generality of heir results would be challenged if the direction of selection in the presence of repentant and forgiving strategies was reversed away from one-shot cooperation. Does this reversal of selection occur?

To see, in the third treatment I introduce a more savvy repentant strategy and a more forgiving cooperative strategy. I replace ALLD with a savvy repentant strategy dubbed “Hesitant Grim” (HGRIM). HGRIM defects on the first round, cooperates on the second, and then plays a trigger strategy where it cooperate until its opponent defects, and continuing to defect thereafter. HGRIM is a fairly simple representative of a repentant strategy. With three memory states, it is comparable in complexity to TF2T (see Appendix D). HGRIM differs from DIMAS in that, while it still cooperates after the first round with repentant and forgiving strategies, it does not cooperate with unrepentant or retaliatory strategies. HGRIM is a simple representative of a large number of strategies with similar properties, such as the nasty, but repentant, version of TFT in the next section.

To simulate the invasion of a novel cooperative strategy, I replace a small fraction (5%) of the initial population with TF2T, with the remainder still playing TFT. TF2T is often a high-performing strategy. In fact, Axelrod submitted it to his own second tournament after determining that it would have won the first, had it been entered (Axelrod, 1984). (TF2T, of course, did not win the second tournament, further highlighting that a strategy’s success hinges on the particular mix of other strategies in the population.)

Having two nice strategies means that selection now acts on two traits in the model. The first is the cue threshold, as in the first two treatments, and the second is the nice strategy (TFT or TF2T) employed with sufficient evidence of a repeated interaction. Therefore, in Treatment 3 each agent has two separate parents from the previous generation (instead of one parent as in Treatments 1 and 2). Each parent is chosen with a probability proportional to their relative payoff. Each agent inherits a cue threshold and a nice strategy from one of its parents drawn randomly and independently for each trait. There is also a 0.1% chance that an agent’s nice strategy will mutate from TFT to TF2T or *vice versa*.

### 2.4. Treatment Four: TFT, TF2T and HTFT

This treatment was inspired by an anonymous reviewer who, concerned about the generalizability of the results of Treatment 3, suggested that a TFT-like strategy would be more plausible than a trigger strategy like HGRIM. Treatment 4 is identical to Treatment 3, except HGRIM is replaced with a strategy called Hesitant Tit-for-Tat, HTFT (McElreath and Boyd, 2007, 170-172). HTFT defects on the first round, cooperates on the second and then plays TFT (see Appendix D). HTFT plays similarly to HGRIM except that, instead of mutual defection with TFT after the first round, HTFT and TFT alternate cooperation and defection. In his evolutionary simulations, Miller (1996) found that both trigger and non-trigger strategies endogenously evolved to play repeated PDs, so it is useful to examine both types of strategies.

## 3. Results

The four treatments described above represent the DKCT model under slightly different mixes of starting strategies. Biasing towards DKCT’s findings, in each treatments agents play nasty strategies with enough evidence that an interaction is one-shot and nice strategies with enough evidence that an interaction is repeated. As in DKCT’s model, the nice strategy played by all of the agents in my first two treatments and the vast majority (95%) of my second two is TFT. Despite this similarity in initial conditions, the direction of selection varies widely between treatments, with one-shot cooperation increasing in many environments in Treatment 1 but decreasing in all environments in Treatments 3 and 4.

Fig. 4 illustrates how introducing repentant and forgiving strategies reverse the direction of selection. Fig. 4A shows that when agents are constrained to ALLD and TFT, similarly to DKCT’s simulations, relatively high probabilities of one-shot cooperation, defined as the expected probability of a randomly selected agent cooperating in a one-shot interaction, can evolve. As shown in Appendix A, one-shot cooperation increases in many of DKCT’s parameter combinations. Specifically, it increases in 608 combinations and decreases in only 142. Given these results, would it be sufficient for a naive agent’s behavior to be evoked merely from its local non-cultural environment before deciding whether to cooperate with a stranger? Or would it be better for the agent to also know something about the locally successful norms?

**Figure 4.**
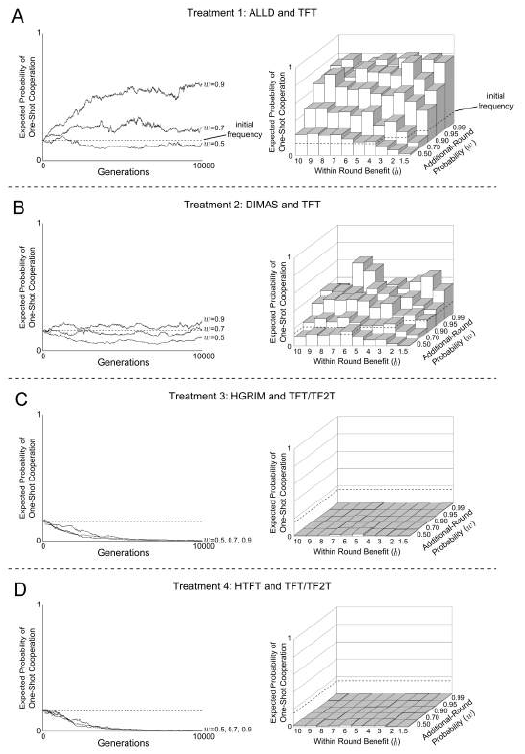
The expected probability of one-shot cooperation substantially decreases with the addition of forgiving and repentant strategies. This figure shows both time-series and final expected probabilities of one-shot cooperation averaged over the last 500 rounds of the simulation for selected parameter combinations (*P* = 0.5, *d* = 2) for Treatment 1 (A), Treatment 2 (B), Treatment 3 (C), and Treatment 4 (D). These parameters match Delton et al. (2011a)’s Fig. 3, though here each agent has ten lifetime partners. Although A shows one-shot cooperation increasing as reported by DKCT, C and D shows that it decreases and virtually disappears when repentant and forgiving strategies are allowed. This general pattern holds across the 750 parameter combinations as shown in Appendix A.

As shown Fig. 4B and Appendix A, merely replacing ALLD with DIMAS, a repentant strategy, reverses the direction of selection away from one-shot cooperation under many parameter combinations, meeting DKCTs criteria for challenging their hypothesis under these combinations. For an agent born into one of these environments it would be unwise to automatically adopt one-shot cooperation or one-shot defection based merely on the local environmental parameters (such as *b*, *P*, and *w*). Depending on the local strategic norms it would sometimes be better to adopt one-shot cooperation (as in Treatment 1) or one-shot defection (as in Treatment 2). A much better strategy would be to observe the local norms and adopt those that were most successful. This is the evolutionary rational for a norm psychology.

In Treatments 3 and 4, where both repentant and forgiving strategies are introduced, selection drives down one-shot cooperation in all conditions (see Fig. 4C and Appendix A). TF2T quickly invades TFT because it captures future cooperation with repentant strategies like HGRIM or HTFT. Eventually, however, strategies that are likely to play HGRIM and HTFT invade because they have higher payoffs than strategies that are more likely to play TF2T. As shown in Appendix A, selection decreases the amount of one-shot cooperation from initial conditions for *all* of DKCT’s 750 parameter combinations in the ten-partner case (though stochastic shocks sometimes overwhelm selection in the one-partner case). An agent born into a society with an evolutionary history of repentance and forgiveness, similar to Treatments 3 or Treatment 4, would not do well if, relying on environmental cues alone, it decided to cooperate on the first round of interaction. It would do better if it identified the prevailing local norms.

Adding repentant and forgiving strategies to the population can reverse the direction of selection away from one-shot cooperation for *all* parameter conditions, clearly meeting Delton et al. (2011b)’s criteria for challenging the generality of their results. Below I show, analytically, that this finding holds even in the case of complete uncertainty.

## 4. The Case of Complete Uncertainty

DKCT credit the high probability of one-shot cooperation to uncertainty over whether an interaction is one-shot or repeated. The strongest case for one-shot cooperation in their model is when uncertainty is maximized, that is when there is no signal indicating whether an agent is in a one-shot or repeated game. This case reduces their model to a standard repeated PD where the probability of transitioning from the first to second round is lower, (1 - *P*)*w*, than the transition probability for every other round, w. In Table 4, I give the conditions where one-shot cooperation is evolutionarily stable and the conditions where it is risk-dominant under the four mixes of strategies described above. Sometimes a population will have more than one equilibrium. Risk dominance indicates that an equilibrium has the largest basin of attraction, implying that the system will be closer to that equilibrium over the long-run average. However, the stability of any particular behavior, as discussed above, is only possible until invaded by a suitable mix of novel strategies. Any equilibrium described here could be invaded by the right mix of novel strategies.

Table 4 gives the general conditions favoring one-shot cooperation decrease when there are repentant and forgiving strategies, which are calculated in Appendix B. We can compare, under each mix of strategies, how many of the 250 combinations of *b*, *w* and *P* that DKCT specify as plausible stabilize one-shot cooperation and in how many is one-shot cooperation risk-dominant. When selection only acts on TFT and ALLD (as in DKCT’s model and my Treatment 1), TFT is stable under a high number, 85%, of the plausible parameter combinations and is also risk-dominant in a majority 65% of them. However when ALLD is replaced with DIMAS, TFT is stable in only 64% of the plausible parameter combinations and is risk-dominant in only 42% of them. (ALLD and DIMAS are always stable in competition with TFT, though their basins of attraction can be very small.) As above, for many environments the successful behavior depends on the existing norms in a society.

**Table 3.** The number of parameter conditions where one-shot cooperation is stable and risk dominant decrease if repentant and forgiving strategies are allowed in the population. When a repentant and forgiving strategy can invade a population with TFT, as in Treatment 3, one-shot cooperation goes to zero.

|  | Treatment 1 | Treatment 2 | Treatment 3 | Treatment 4 |
| --- | --- | --- | --- | --- |
| One-Shot Coop.<br>is Stable | $\frac{c}{b} < \frac{w(1-P)}{1-wP}$ | $\frac{c}{b} < w(1-P)$ | — | — |
| # Parameter<br>Combinations | 237:250<br>(85%) | 160:250<br>(64%) | 0:250<br>(0%) | 0:250<br>(0%) |
| One-Shot Coop.<br>Risk-Dominant | $\frac{c}{b} < \frac{w(1-P)}{2(1-wP)}$ | $\frac{c}{b} < \frac{1}{2}w(1-P)$ | — | — |
| # Parameters<br>Combinations | 163:250<br>(65%) | 106:250<br>(42%) | 0:250<br>(0%) | 0:250<br>(0%) |

Finally, in a population with HGRIM, TFT and TF2T (as in Treatment 3) or with HTFT, TFT and TF2T (as in Treatment 4) one-shot cooperation in the repeated PD is *never* stable or risk-dominant under any possible combinations of *w*, *P*, or *b*, as I show in Appendix B. We would expect, as I found in simulation, that one-shot cooperation would decrease in any such a population and eventually disappear. That one-shot cooperation is so readily replaced by one-shot defection when a population includes repentant and forgiving strategies, even under conditions of complete uncertainty, is a problem for any hypothesis premised on reciprocity under uncertainty leading inevitably to cooperative equilibria. That there is such variation in the direction of selection for the same environments across conditions is also a problem for any hypothesis premised on the idea that rules for social exchange need only be evoked from the non-cultural environment.

## 5. Generalizability of Results

Because, in an infinitely repeated game, the space of potential strategies is infinitely large, I agree with Delton et al. (2011b) that it is useful to choose representative strategies to illustrate key trade-offs that must be made by selection. However, some may wonder about these strategies’ generalizability. The above treatments include strategies that are simple representatives of a vast universe of repentant strategies (like DIMAS, HGRIM, HTFT) and a vast universe of nice, but retaliatory, strategies (like TFT and TF2T). For example, in Treatment 2 DIMAS is equivalent in payoffs to, say, Hesitant Tit-for-Two-Tats, a strategy like HTFT, but playing TF2T instead of TFT. DIMAS is the simplest repentant strategy in terms of computational complexity (Miller, 1996), but evolution could generate a large set of similar strategies. In Treatments 3 and 4, TF2T could be replaced by, say, a strategy that always cooperates or TF3T with identical results. HGRIM and HTFT are simple representatives of an infinite number of hesitant strategies that would act similarly under selection. They are, in fact, *less* sophisticated versions of the type of ”apologetic” strategies that evolved endogenously in Axelrod (1997)’s simulations. For any simple strategy, there is an infinite set of strategies of more computational complexity that will behave similarly or identically in the same environmental and cultural conditions. As discussed in Section 1.2, selection can always find a combination of these strategies that will dislodge existing behavioral equilibria.

Even so, some might question whether repentant strategies are appropriate strategies to be considered at all. Delton et al. (2011b)’s reply to McNally and Tanner (2011) states that hesitant strategies “introduce aspects of cooperation beyond the empirical issue we were addressing: the human propensity to be generous with a novel partner on the first move in a situation that appears to be one-shot. In such situations, a hesitant strategy - unlike humans - would not be generous on the first move and could thus not plausibly model the actual behavior we sought to understand.” This argument is problematic. If one’s goal is to demonstrate how one-shot cooperation emerges, through selection, from a set of possible alternative behaviors, it is inappropriate to assume, *a priori*, that those alternative behaviors cannot exist. If a hypothesis cannot account for selective elimination of competing strategies, that hypothesis should either be modified or replaced. This, as I describe in the next section, is where the norm psychology hypothesis comes in.

## 6. Discussion

I have demonstrated that repeated interaction, even under uncertainty, does not necessarily favor one-shot cooperation. However, my aim is *not* to show that one-shot defection is a more likely outcome than one-shot cooperation. We know, from the results described in Section 1.2, that the strategies dominating Treatments 3 and 4 could, themselves, be eventually invaded by the right mix of other strategies. My claim is that many of these trajectories are possible and, even under the same environmental conditions, the direction of selection depends on a particular group’s idiosyncratic history of invasion.

This paper compares the logical underpinnings of two competing hypotheses seeking to explain the prevalence of cooperation in one-shot laboratory experiments. The mismatch hypothesis, which is based on the premise that repeated interaction leads to a bias towards one-shot cooperation, found support in a recent model. However, the increase of one-shot cooperation in the model was due to agents’ evolutionary possibilities being artificially constrained to only two strategies, TFT and ALLD. The model’s authors suggest that the model would lose its generality if the addition of repentant and forgiving strategies reversed selection away from one-shot cooperation (Delton et al., 2011b). Here I demonstrate that the addition of forgiving and repentant strategies reverses the direction of selection for all plausible environmental conditions. In total, these models suggest that societies with different evolutionary histories will have very different levels of one-shot cooperation. This finding is consistent with the conditions for the norm psychology hypothesis which is premised on the idea that humans evolved, through a process of gene-culture coevolution, to flexibly learn and adopt the particular norms of their particular society.

Both DKCT’s and my simulations are stylized models of selection. Selection is a general process of variation and differential transmission that is, itself, substrate neutral (Dennett, 1995). In translating the stylized model to real world analogs, DKCT suppose that cue summaries result from genetic selection on genetically-transmitted information and I suppose that they result from cultural selection operating on socially-transmitted information. My translation is consistent with the norm psychology hypothesis, which supposes that once humans evolved to rely to a large extent on social learning, cultural traits became subject to selection-like forces, especially if cultural traits of more successful individuals are more likely to spread. Because cultural traits, unlike genetic traits, can be transmitted horizontally to many conspecifics within a single lifetime, cultural evolution can occur much more rapidly than genetic evolution. Therefore, different groups of humans, even with similar genetic inheritance, can quickly establish very different local norms. Norms arising within a group are influenced by the local environmental conditions, but groups in nearly identical environments can have very different norms. One reason, as in the model above, is because patterns of repeated interaction allow *any* behavior to become an equilibrium, with the caveat that all of these equilibria are unstable.

However, in the norm psychology hypothesis, at least two forces, conformity and punishment, can stabilize norms. If humans have a predisposition towards adopting the more common behavior in their group (which will be adaptive under many conditions (Boyd and Richerson, 1985; Henrich and Boyd, 1998, Ch. 11)) or norms are enforced by punishment (Boyd and Richerson, 1992), norms will be slower to change. While costly punishment is subject to so-called “second order” effects, because punishment is itself a costly normative behavior, conformity and punishment operate together to overcome these second-order effects (Henrich and Boyd, 2001). That cultural transmission is often conformist, but genetic transmission is not, is one reason cultural inheritance is thought to be a more likely route to costly cooperation than pure genetic inheritance (Bell, 2010).

How does the norm psychology hypothesis explain that cooperation in economic experiments is common in many, but not all, societies? The norm psychology hypothesis has a close cousin, the “cultural group selection hypothesis” which examines the consequences of the social learning mechanisms underlying norm psychology in group-structured populations. The general hypothesis, famously proposed by Charles Darwin (1873), is that when groups of humans are in competition, groups where individuals are more cooperative will tend to out-compete groups where cooperation is rare.

For example, consider groups undergoing cultural evolution under complete uncertainty in a strategy space similar to that in Treatment 2, where both TFT and DIMAS are stable equilibria and neither is risk-dominant, as occurs when *w* = 0.5, *P* = 0.5 and *b* = 8. The norms of these groups, evolving in isolation, should be just as likely to converge to a TFT-playing equilibrium as to a DIMAS-playing equilibrium. Variation between groups is maintained in the face of migration because a DIMAS-playing agent would fair poorly in a TFT-playing group and *vice versa*. It would be best to adopt the norms of the new group rather than keep the norms of the old group (an option available to cultural, but not genetic, inheritance). However, a population of TFT-playing agents would have higher overall payoffs than DIMAS-playing agents. If these groups come into competition, TFT-playing groups should out-compete DIMAS-playing groups and norms for one-shot cooperation would, on average, spread. Of course, if evolution has access to the entire strategy space we should see a greater diversity of normative equilibria than the pure equilibria of these two strategies.

This process of “equilibrium selection” is generalizable to any case where there is variation between competing groups and the relative success of the groups depend on this variation (Boyd and Richerson, 1990). Modeling and empirical measurements suggest that, because cultural evolution is more rapid and subject to conformity, between-group variation is likely to be greater when behavior is transmitted socially than genetically. This is because when a norm psychology induces humans to adopt their group’s traits, this further reinforces any existing equilibrium and drives down within-group variance relative to between-group variance (Henrich, 2004; Bell, 2010).

This suggests how reciprocity under uncertainty might fit into the larger picture of human evolution: it is one of many mechanisms for generating between-group variation in cultural norms. Because the norms of groups can evolve in radically different directions, even in similar environments, it was not good enough for one’s behavior to simply be “evoked” from the environment. Instead, humans likely evolved to identify and adopt the successful norms for their particular group. The behavior of an individual in an economic experiment reflects their norms and the distribution of individuals’ behavior within a group is expected to co-vary with the group’s distribution of norms. When groups of individuals with different norms for social exchange come into competition, the group with the more cooperative norms will, all else equal, out-compete groups with less cooperative norms. Thus, as Darwin (1873) writes, “the social and moral qualities would tend slowly to advance and be diffused throughout the world.”

## Acknowledgments

I thank Peter Richerson, Richard McElreath, Kyle Joyce, Emily Peffer, Katie Demps, Kari Schroeder, Bruce Winterhalder, members of the UC Davis Cultural Evolution and Human Behavioral Ecology Labs and three anonymous reviewers for comments on earlier drafts of this paper and Andrew Delton and Max Krasnow for help in recreating their original model. This work was funded, in part, by a block grant from the University of California, Davis, Graduate Group in Ecology and partly preformed at NIMBioS, sponsored by the NSF, U.S. Department of Homeland Security, and U.S. Department of Agriculture through NSF Awards EF-0832858 and DBI-1300426, with additional support from the University of Tennesse

## Appendix A Probability of One-Shot Cooperation for All Parameters and Treatments

**Figure A.1.**
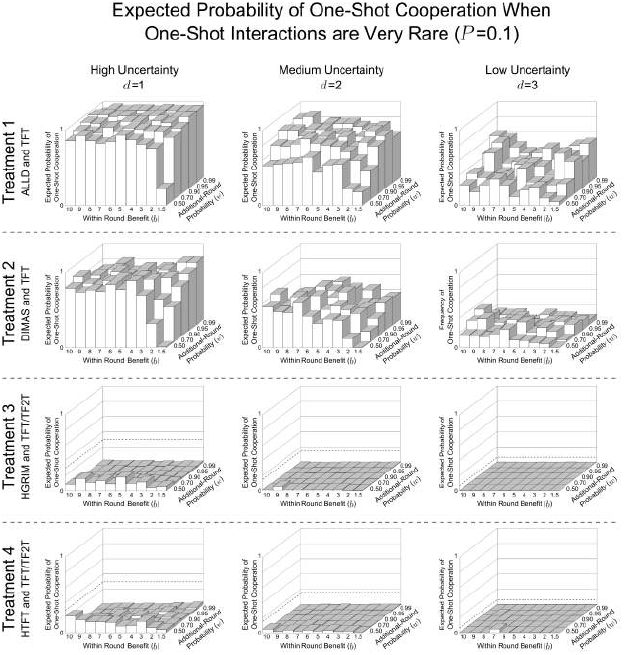
Repentant and forgiving strategies decrease one-shot cooperation when one-shot games are very rare (*P* = 0.1) and agents have 10 partners. These show the expected probability of one-shot cooperation averaged over the last 500 generations of the 10,000 generation simulation for all values of *d*, *b*, and *w*. These are the same parameter combinations reported by Delton et al. (2011a). Treatment 1, where agents, as in DKCT play only TFT or ALLD has the highest probability of one-shot cooperation. Treatment 2, where ALLD is replaced by a repentant strategy, DIMAS, has less one-shot cooperation. In Treatment 3, where a savvy repentant strategy, HGRIM, competes with TFT and TF2T, one-shot cooperation virtually disappears. And Treatment 4, where HGRIM is replaced by HTFT. This highlights the variability in outcomes for the same environmental parameters.

**Figure A.2.**
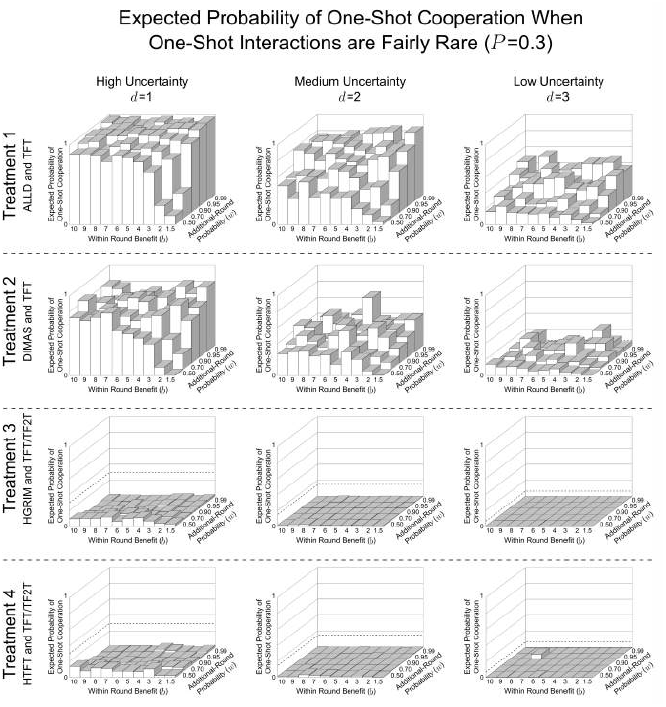
Repentant and forgiving strategies decrease one-shot cooperation when one-shot games are fairly rare (*P* = 0.3) and agents have 10 partners. These show the expected probability of one-shot cooperation averaged over the last 500 generations of the 10,000 generation simulation for all values of *d*, *b*, and *w*. These are the same parameter combinations reported by Delton et al. (2011a). Treatment 1, where agents, as in DKCT play only TFT or ALLD has the highest probability of one-shot cooperation. Treatment 2, where ALLD is replaced by a repentant strategy, DIMAS, has less one-shot cooperation. In Treatment 3, where a savvy repentant strategy, HGRIM, competes with TFT and TF2T, one-shot cooperation virtually disappears. And Treatment 4, where HGRIM is replaced by HTFT. This highlights the variability in outcomes for the same environmental parameters.

**Figure A.3.**
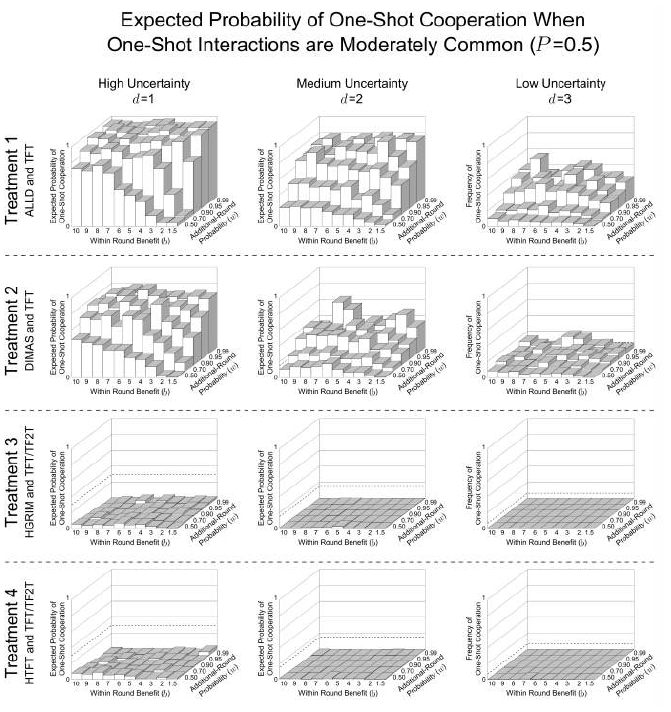
Repentant and forgiving strategies decrease one-shot cooperation when one-shot games are moderately rare (*P* = 0.5) and agents have 10 partners. These show the expected probability of one-shot cooperation averaged over the last 500 generations of the 10,000 generation simulation for all values of d, b, and w. These are the same parameter combinations reported by Delton et al. (2011a). Treatment 1, where agents, as in DKCT play only TFT or ALLD has the highest probability of one-shot cooperation. Treatment 2, where ALLD is replaced by a repentant strategy, DIMAS, has less one-shot cooperation. In Treatment 3, where a savvy repentant strategy, HGRIM, competes with TFT and TF2T, one-shot cooperation virtually disappears. And Treatment 4, where HGRIM is replaced by HTFT. This highlights the variability in outcomes for the same environmental parameters.

**Figure A.4.**
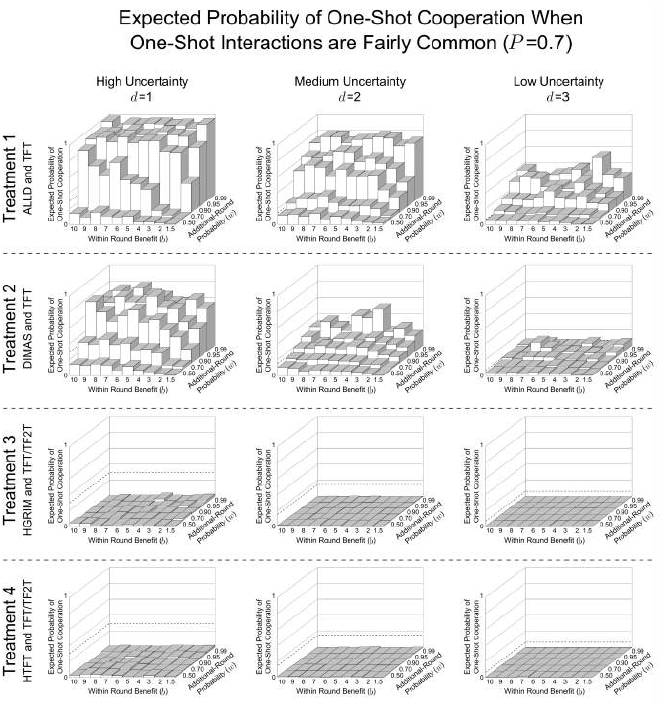
Repentant and forgiving strategies decrease one-shot cooperation when one-shot games are fairly common (*P* = 0.7) and agents have 10 partners. These show the expected probability of one-shot cooperation averaged over the last 500 generations of the 10,000 generation simulation for all values of *d*, *b*, and *w*. These are the same parameter combinations reported by Delton et al. (2011a). Treatment 1, where agents, as in DKCT play only TFT or ALLD has the highest probability of one-shot cooperation. Treatment 2, where ALLD is replaced by a repentant strategy, DIMAS, has less one-shot cooperation. In Treatment 3, where a savvy repentant strategy, HGRIM, competes with TFT and TF2T, one-shot cooperation virtually disappears. And Treatment 4, where HGRIM is replaced by HTFT. This highlights the variability in outcomes for the same environmental parameters.

**Figure A.5.**
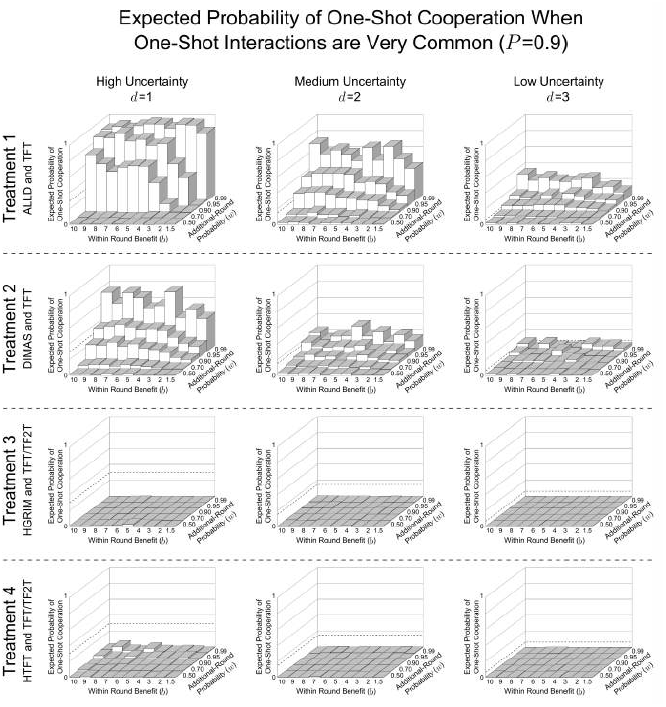
Repentant and forgiving strategies decrease one-shot cooperation when one-shot games are very common (*P* = 0.9) and agents have 10 partners. These show the expected probability of one-shot cooperation averaged over the last 500 generations of the 10,000 generation simulation for all values of *d*, *b*, and *w*. These are the same parameter combinations reported by Delton et al. (2011a). Treatment 1, where agents, as in DKCT play only TFT or ALLD has the highest probability of one-shot cooperation. Treatment 2, where ALLD is replaced by a repentant strategy, DIMAS, has less one-shot cooperation. In Treatment 3, where a savvy repentant strategy, HGRIM, competes with TFT and TF2T, one-shot cooperation virtually disappears. And Treatment 4, where HGRIM is replaced by HTFT. This highlights the variability in outcomes for the same environmental parameters.

**Figure A.6.**
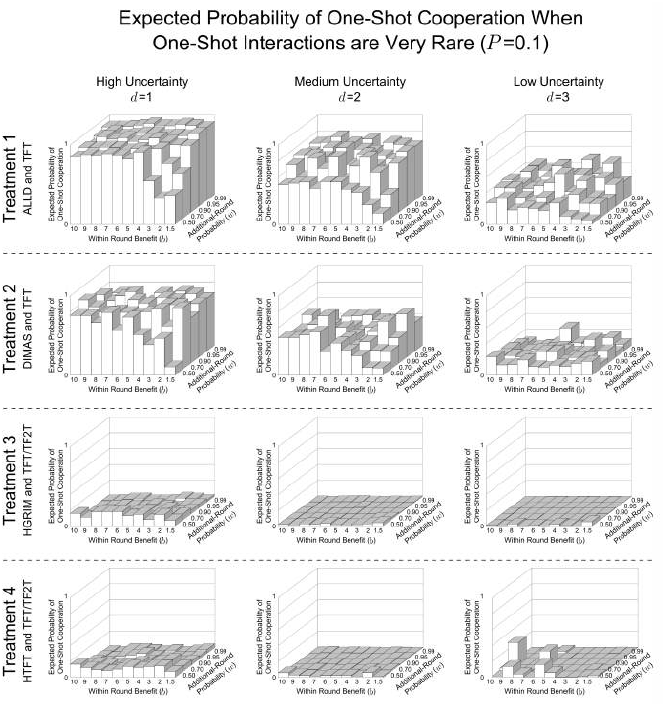
Repentant and forgiving strategies decrease one-shot cooperation when one-shot games are very rare (*P* = 0.1) and agents have one partner. These show the expected probability of one-shot cooperation averaged over the last 500 generations of the 10,000 generation simulation for all values of *d*, *b*, and *w*. These are the same parameter combinations reported by Delton et al. (2011a). Treatment 1, where agents, as in DKCT play only TFT or ALLD has the highest probability of one-shot cooperation. Treatment 2, where ALLD is replaced by a repentant strategy, DIMAS, has less one-shot cooperation. In Treatment 3, where a savvy repentant strategy, HGRIM, competes with TFT and TF2T, one-shot cooperation virtually disappears. And Treatment 4, where HGRIM is replaced by HTFT. This highlights the variability in outcomes for the same environmental parameters.

**Figure A.7.**
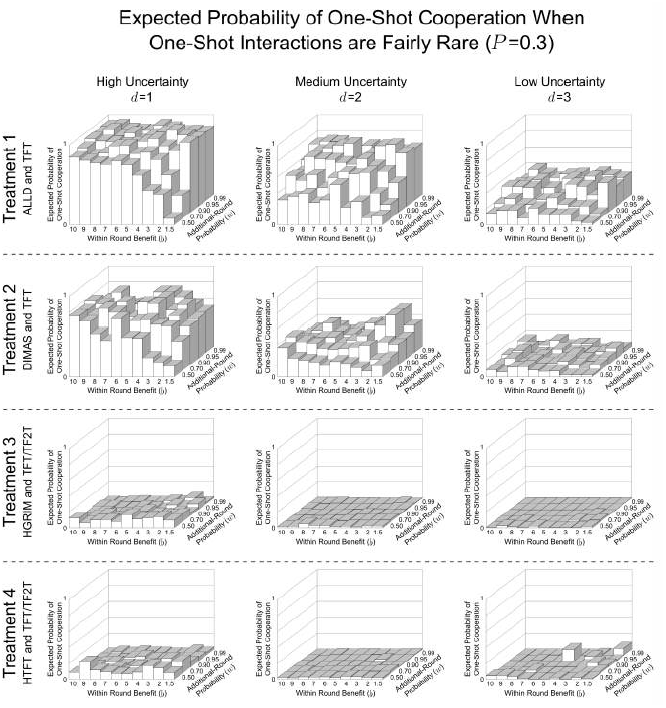
Repentant and forgiving strategies decrease one-shot cooperation when one-shot games are fairly rare (*P* = 0.3) and agents have one partner. These show the expected probability of one-shot cooperation averaged over the last 500 generations of the 10,000 generation simulation for all values of *d*, *b*, and *w*. These are the same parameter combinations reported by Delton et al. (2011a). Treatment 1, where agents, as in DKCT play only TFT or ALLD has the highest probability of one-shot cooperation. Treatment 2, where ALLD is replaced by a repentant strategy, DIMAS, has less one-shot cooperation. In Treatment 3, where a savvy repentant strategy, HGRIM, competes with TFT and TF2T, one-shot cooperation virtually disappears. And Treatment 4, where HGRIM is replaced by HTFT. This highlights the variability in outcomes for the same environmental parameters.

**Figure A.8.**
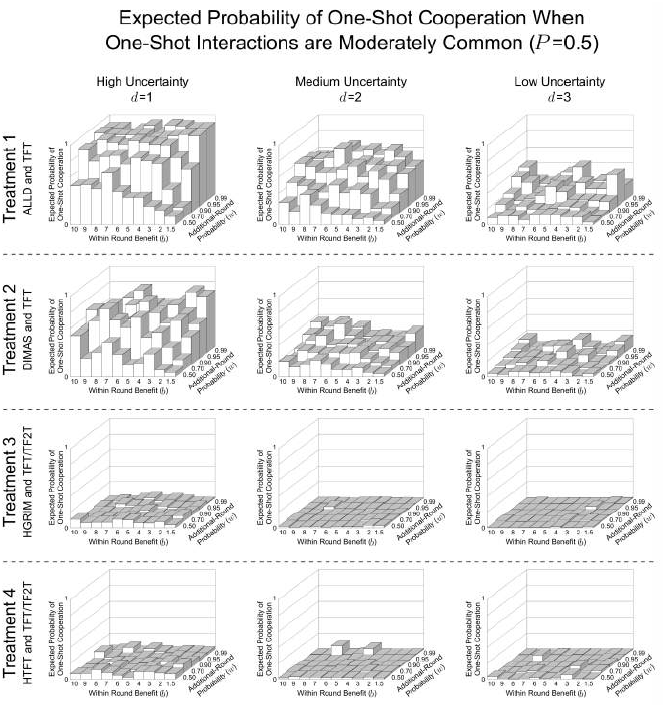
Repentant and forgiving strategies decrease one-shot cooperation when one-shot games are moderately rare (*P* = 0.5) and agents have one partner. These show the expected probability of one-shot cooperation averaged over the last 500 generations of the 10,000 generation simulation for all values of *d*, *b*, and *w*. These are the same parameter combinations reported by Delton et al. (2011a). Treatment 1, where agents, as in DKCT play only TFT or ALLD has the highest probability of one-shot cooperation. Treatment 2, where ALLD is replaced by a repentant strategy, DIMAS, has less one-shot cooperation. In Treatment 3, where a savvy repentant strategy, HGRIM, competes with TFT and TF2T, one-shot cooperation virtually disappears. And Treatment 4, where HGRIM is replaced by HTFT. This highlights the variability in outcomes for the same environmental parameters.

**Figure A.9.**
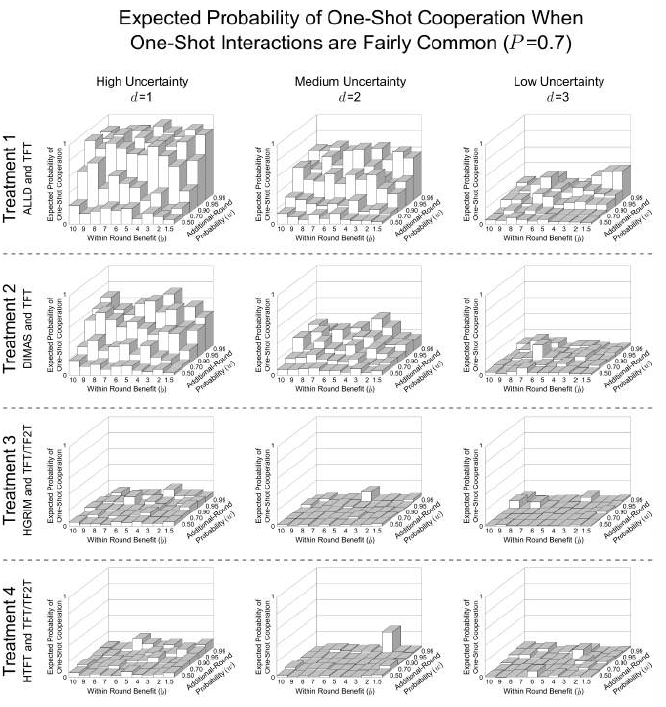
Repentant and forgiving strategies decrease one-shot cooperation when one-shot games are fairly common (*P* = 0.7) and agents have one partner. These show the expected probability of one-shot cooperation averaged over the last 500 generations of the 10,000 generation simulation for all values of *d*, *b*, and *w*. These are the same parameter combinations reported by Delton et al. (2011a). Treatment 1, where agents, as in DKCT play only TFT or ALLD has the highest probability of one-shot cooperation. Treatment 2, where ALLD is replaced by a repentant strategy, DIMAS, has less one-shot cooperation. In Treatment 3, where a savvy repentant strategy, HGRIM, competes with TFT and TF2T, one-shot cooperation virtually disappears. And Treatment 4, where HGRIM is replaced by HTFT. This highlights the variability in outcomes for the same environmental parameters. Some populations show increased one-shot cooperation. This is due to stochastic shocks as explained in Appendix C.

**Figure A.10.**
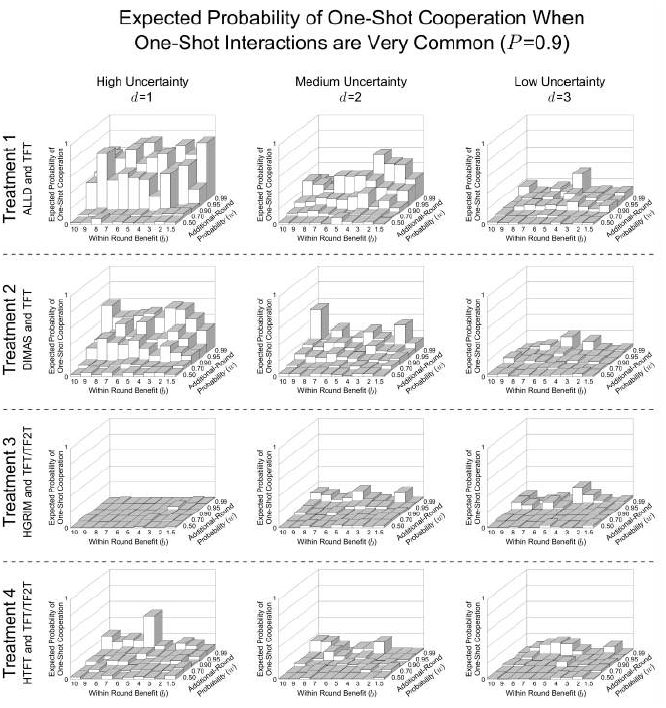
Repentant and forgiving strategies decrease one-shot cooperation when one-shot games are very common (*P* = 0.9) and agents have one partner. These show the expected probability of one-shot cooperation averaged over the last 500 generations of the 10,000 generation simulation for all values of *d*, *b*, and *w*. These are the same parameter combinations reported by Delton et al. (2011a). Treatment 1, where agents, as in DKCT play only TFT or ALLD has the highest probability of one-shot cooperation. Treatment 2, where ALLD is replaced by a repentant strategy, DIMAS, has less one-shot cooperation. In Treatment 3, where a savvy repentant strategy, HGRIM, competes with TFT and TF2T, one-shot cooperation virtually disappears. And Treatment 4, where HGRIM is replaced by HTFT. This highlights the variability in outcomes for the same environmental parameters. Some populations show increased one-shot cooperation. This is due to stochastic shocks as explained in Appendix C.

## Appendix B Complete Uncertainty Calculations

The best-case scenario for the evolution of one-shot cooperation in the DKCT model is under the condition of complete uncertainty, where there is no signal concerning whether a game is one-shot or repeated. Under complete uncertainty, DKCT’s model reduces to a standard repeated Prisoner’s Dilemma where the probability of transitioning from the first to second round is (1 − *P*)*w*, which is lower than the transition probability for every other round, *w*. Fig. B.1 gives the expected payoffs for the strategy combinations in all four treatments under complete uncertainty. In this appendix, I show the calculations used to derive the stability and risk-dominance conditions in Table 4.

**Figure B.1.**
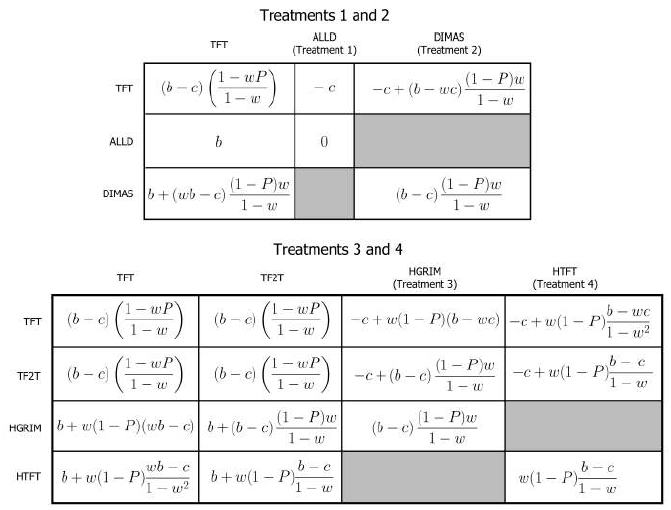
Expected payoffs for strategies in the four treatments when agents are completely uncertain whether a game is one-shot or repeated. *b* is the benefits to cooperation, *c* is the cost of cooperating, *P* is the probability that an interaction is one-shot, and w is the probability that, after each round in a repeated game, there is another round. TFT occurs in all treatments. ALLD occurs in Treatment 1. DIMAS occurs in Treatment 2. HGRIM occurs in Treatment 3. HTFT occurs in Treatment 4. TF2T occurs in Treatments 3 and 4.

### Appendix B.1. Treatment 1: When is TFT stable against ALLD?

TFT is stable against ALLD when the payoff to TFT given TFT is greater than the payoff to ALLD given TFT:

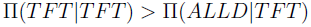

Substituting from Fig. B.1 and simplifying yields the condition where TFT, and thus one-shot cooperation is stable:

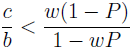

### Appendix B.2. Treatment 2: When is TFT stable against DIMAS?

TFT is stable against DIMAS when the payoff to TFT given TFT is greater than the payoff to DIMAS given TFT:

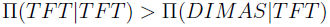

Substituting from Fig. B.1 and simplifying yields the condition where TFT, and thus one-shot cooperation is stable:

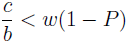

### Appendix B.3. Treatment 3: When is TFT stable against direct invasion by HGRIM?

TFT is stable against HGRIM when the payoff to TFT given TFT is greater than the payoff to HGRIM given TFT:

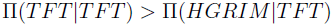

Substituting and Simplifying:

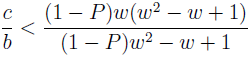

### Appendix B.4. When is TFT stable against indirect invasion by HGRIM via TF2T?

One of two conditions must be met for TFT to be stable against indirect invasion by HGRIM via TF2T. First the payoff to TFT given HGRIM could be greater than the payoff to TF2T given HGRIM:

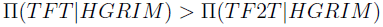

Substituting from Fig. B.1 and simplifying yields the condition where TFT, and thus one-shot cooperation is stable:

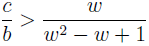

Second the payoff to TF2T given HGRIM may be greater than the payoff to TF2T given TF2T.

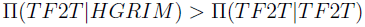

By inspection of Fig. B.1 this can never be true because these payoffs are equivalent in all rounds except the first where TF2T gains a benefit against itself, but not against HGRIM..

### Appendix B.5. *When is TFT stable against both* direct *and* indirect *invasion by HGRIM?*

From Appendix B.3 and Appendix B.4, the condition where TFT is stable against both direct and indirect invasion is:

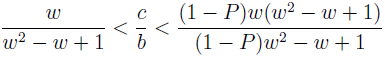

It is easy to show that 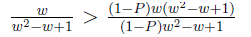 over all possible values of *P*, *w*, and 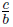, therefore TFT is never stable against co-invasion by TFT and HGRIM.

### Appendix B.6. *Treatment 4: When is TFT stable against* direct *invasion by HTFT?*

The payoff matrix for this HTFT, TFT and TF2T under complete uncertainty is shown in Fig. B.1.

TFT is stable against direct invasion by HTFT when the payoff to TFT given TFT is greater than the payoff to HTFT given TFT:

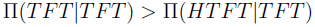

Substituting from the matrix in Fig. B.1:

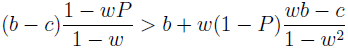

After algebraic manipulation, TFT is stable against direct invasion by HTFT when:

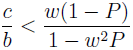

### Appendix B.7. When is TFT stable against indirect invasion by HTFT via TF2T?

Either of two conditions must be met for TFT to be stable against indirect invasion by HTFT via TF2T. First, the payoff to TFT given HTFT could be higher than the payoff to TF2T given HTFT:

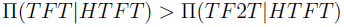

Substituting from the matrix in Fig. B.1:

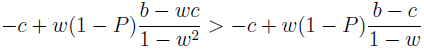

After algebraic manipulation, TFT is stable to invasion by TF2T in the presence of HTFT when:

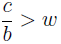

Second, the payoff to TF2T given TF2T could be greater than the payoff to HTFT given TF2T.

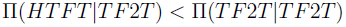

Substituting from the matrix in Fig. B.1:

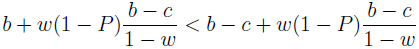

After algebraic manipulation, TF2T invades TFT in the presence of HTFT when:

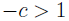

This is never true, so TFT is only stable to invasion by HTFT in the presence of TF2T when:

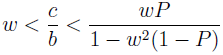

It is easy to show that 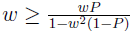 for all *P* ∈ [0:1] and all *w* ∈ [0:1). Therefore TFT can always be invaded by HTFT in the presence of TF2T as long as the number of rounds is not infinite.

## Appendix C Effect of Increasing Partners from One to Ten

The analytic analysis presented in Table 4 indicates that, in Treatments 3 and 4, one-shot cooperation should decrease under all parameter conditions. However, Figures A.9 and A.10 show some instances of increased one-shot cooperation in these treatments, especially when the probability of a one-shot interaction (*P*) and the probability of an additional round given a repeated interaction (*w*) are high. It turns out that this cooperation is maintained as a by-product of a large amount of stochasticity in agents’ payoffs due to a highly skewed distribution in rounds played. This effect was not apparent in DKCT’s original simulations because, when agents can only play TFT and ALLD, the amount of repeated interaction only matters for agents’ payoffs when both are playing TFT. But when there are patterns of repentant and forgiving strategies, the distribution of rounds in repeated games becomes more important (Fig. 3).

Fig. C.1A shows the probability density function (generated numerically from 10 million samples) for the number rounds per partner an agent plays when agents have only one partner and P and w are high (*P* = 0.9 and *w* = 0.99). This distribution has a very long tail. In expectation, more than 90% of agents agents play only one round in their lifetime, but one dyad per generation is expected to play more than 318 rounds - over 3.5 times as many rounds as the bottom 90% of dyads *combined*. If this dyad happens to be a pair of agents with under-performing, but cooperative strategies, like TFT in Treatments 3 and 4, the strategy would increase dramatically in the population, not because it is a better strategy, but because it was lucky enough to receive a large number of rounds. Taking the whole 10,000 generation simulation, where one dyad is expected to play more than 1225 rounds, into account makes the potential for a stochastic shock even more apparent.

**Figure C.1.**
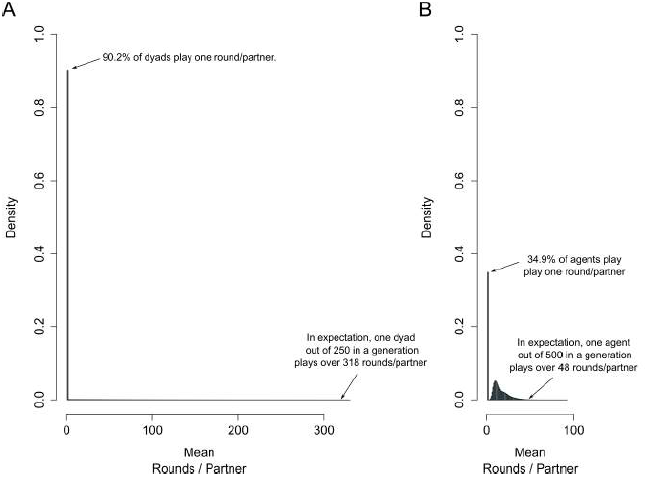
Increasing the number of partners per agent decreases the magnitude of outliers in the number of rounds played per partner. **A** shows the probability density function for number of rounds played by dyads under the conditions *P* = 0.1 and *w* = 0.99. B shows the probability density function for number rounds of played per partner, if each agent has ten partners, under the same conditions

This effect would be interesting if it reflected real world distributions in patterns of interaction. However, this extreme skew is an artifact of agents having only one interaction partner in their lifetime. The amount of skew (as one would expect from the Central Limit Theorem) is drastically reduced when agents interact with multiple other agents. Fig. C.1B shows the probability density function under the same parameters when when agents play with ten partners. The distribution of the number of rounds, normalized to rounds per partner, is much less skewed. Here only 34.9% of agents play one round per partner. In a given generation one agent is expected to play greater than 48 rounds/partner. Out of 10 million samples used to generate Fig. C.1B, *no* agents played over 92 rounds per partner.

Due to the Central Limit Theorem, the expectation of number of rounds per partner in each distribution is the same (i.e., 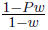), but playing with multiple partners is more realistic, less skewed and limits the effect of stochastic shocks, and still maintains the logical structure of the game itself.

## Appendix D Finite State Machine Representations of Strategies

Strategies from all four treatments are represented in Fig. D.1 as Finite State Automata. Initial plays of the strategy are represented by the state in the double circle (nasty strategies start with Defect and nice strategies start with Cooperate). Transition rules are represented by arrows. For example, HGRIM starts with Defect, transitions to Cooperate, and stays at Cooperate until its opponent defects. After defection by an opponent, DIMAS transitions to Defect where it stays. HTFT is similar, except that once it reaches Defect, it can transition back to Cooperate if its opponent cooperates.

**Figure D.1.**
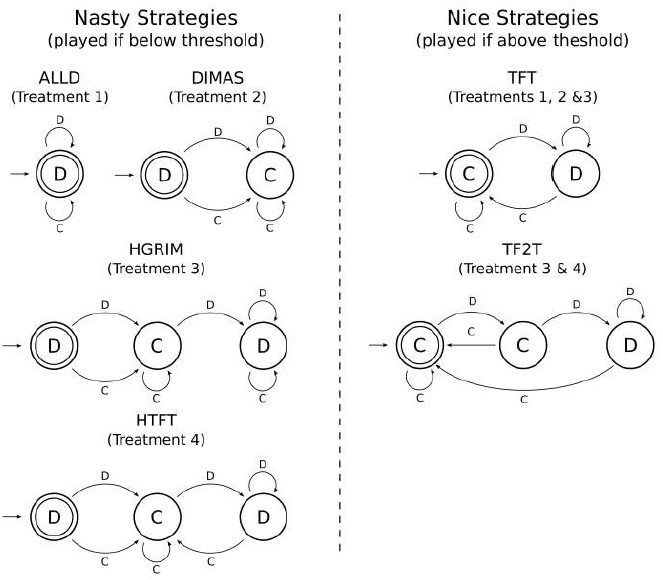
The strategies included in all four treatments represented as Moore Machines, a class of Finite State Automata. The number of states in the minimal FSA is a measure of the complexity of a strategy. ALLD is a one-state strategy. TFT and DIMAS are two-state strategies. TF2T and HGRIM are three-state strategies.

## Footnotes

1 DKCT report similar results from a version of the model with the same game structure and strategy space, but where agents have more complicated cognitive architecture. Since the logic of payoff-based selection applies to any cognitive architecture with sufficient flexibility, for tractability, I focus on the simpler model.

